# PepPCBench is a Comprehensive Benchmark for Protein-Peptide Complex Structure Prediction with AlphaFold3

**DOI:** 10.1101/2025.04.08.647699

**Authors:** Silong Zhai, Huifeng Zhao, Jike Wang, Shaolong Lin, Tiantao Liu, Dejun Jiang, Huanxiang Liu, Yu Kang, Xiaojun Yao, Tingjun Hou

**Affiliations:** Faculty of Applied Science, Macao Polytechnic University, Macao 999078; College of Pharmaceutical Sciences, Zhejiang University, Hangzhou 310058, Zhejiang, China; XiangYa College of Pharmaceutical Sciences Central South University

**Author notes:** These authors contributed equally to this work (Silong Zhai, Huifeng Zhao).

**Keywords:** Protein-peptide complex, Structure prediction, AlphaFold3, Benchmark, Peptide-based therapeutics

## Abstract

Peptides mediate up to 40% of protein-protein interactions (PPIs), offering high specificity and the ability to target binding sites inaccessible to small molecules, making them promising candidates for drug development. Accurate modeling of protein-peptide complexes is crucial for understanding fundamental biological processes and for advancing peptide-based drug design. However, due to the high conformational flexibility of peptides, predicting their interactions with proteins remains a significant challenge. Recent advancements in artificial intelligence (AI), exemplified by all-atom protein folding neural networks (PFNNs) such as AlphaFold3 (AF3), have expanded predictive capabilities beyond proteins to encompass protein-peptide complexes. Nevertheless, existing evaluations of these methods are often limited in scope and lack systematic, multi-dimensional, and fair comparisons of PFNN performance in protein-peptide complex prediction. Here, we introduce PepPCBench, a comprehensive benchmark framework specifically developed to evaluate AF3’s capabilities in predicting protein-peptide complexes. This study utilizes a carefully curated dataset named PepPCSet, which is excluded from the AF3’s training or validation sets. This dataset includes 261 protein-peptide complexes with peptide lengths ranging from 5 to 30 residues. Our benchmark results indicate that AF3 outperforms other PFNNs in prediction accuracy and structural validation. However, its performance remains insufficient for practical peptide drug discovery, indicating room for improvement. It is expected that PepPCBench can provide some valuable insights into the enhancement of protein-peptide complex structure prediction and the development of peptide-based therapeutics. The dataset and pipeline protocols are available at https://github.com/zhaisilong/PepPCBench.

## 1 Introduction

Protein-protein interactions (PPIs) are crucial for key biological processes, including signal transduction and molecular transport, making them attractive drug targets.^1–3^ Unlike antibodies, which struggle with membrane permeability, peptides offer a unique advantage: they combine high specificity and strong binding affinity with ease of synthesis, low toxicity, and minimal immunogenicity.^4^ These properties make peptides ideal for targeting previously “undruggable” PPI interfaces, opening new possibilities for drug development. To date, over 100 peptide-based drugs have been approved by the Food and Drug Administration (FDA), and many more are in the pipeline.^5^ However, designing peptides to modulate PPIs remains a significant challenge due to their inherent conformational flexibility, which complicates accurate binding prediction.^6^ Traditional structure determination techniques, such as X-ray crystallography, are resource-intensive and time-consuming.^7^ Meanwhile, computational approaches, particularly blind protein-peptide docking, struggle with the absence of known peptide structures, in contrast to classical domain-domain docking, where the structures of free individual domains are well-established.^8^ To overcome these limitations, the development of more sophisticated modeling strategies is essential to better predict and capture protein-peptide interactions (PpIs), thus advancing peptide-based therapeutics.

Recent breakthroughs in artificial intelligence (AI)-driven protein structure prediction have revolutionized structural biology.^9^ Among these innovations, the AlphaFold^9,10^ (AF) series has emerged as a pivotal advancement in solving the protein folding problem. In the context of peptide-related research, AF offers a “co-folding” strategy for addressing the global peptide docking challenge by conceptualizing peptide binding as the final step of protein folding, where the peptide completes the receptor surface like a missing piece.^11^ Some studies have shown that docking performance with AF tools typically outperforms traditional methods.^8,12,13^ The latest version, AlphaFold3^14^ (AF3), marks a significant improvement, as it extends its capabilities to model chemical modifications, such as non-standard amino acids and unconventional cyclic structures. This is an important advancement, as existing methods struggle to handle such complexities and often come with high computational costs and limited accuracy. With AF3, the ability to model a wider variety of biomolecules with improved prediction accuracy has the potential to reshape the landscape of structural biology, especially for complex protein-peptide interactions.

However, despite these advances, the application of AF3 to protein-peptide modeling remains an open question. While AF3 addresses many challenges, it does not fully resolve the difficulties inherent in predicting protein-peptide interactions in real-world drug discovery scenarios. Challenges such as peptide flexibility, binding specificity, and conformational changes still need to be addressed.^15^ These limitations highlight the need for further refinement before AF3 can be seamlessly integrated into peptide-based drug design workflows. To address these gaps, comprehensive benchmarks for protein-peptide complex structure prediction are needed to assess both the upper and lower performance limits of AF3. Additionally, further development of complementary tools and strategies is necessary to improve these aspects and mitigate their negative impact, tailored to specific tasks.

Previous benchmarks of protein-peptide complexes have primarily focused on traditional docking methods and molecular simulations.^16,17^ However, there has been a lack of benchmarks specifically assessing folding neural networks for protein-peptide interactions. A recent notable study by Manshour et al.^18^ tried to address this gap by compiling a dataset of 60 protein-peptide complexes from the Protein Data Bank^19^ (PDB) and evaluating AlphaFold-Multimer^20^ (AFM), AF3, and ColabFold^21^ (CF) with scoring functions for predicting protein-peptide complex structures. Building upon their efforts, we aim to expand the scope and depth of peptide-based research, enhancing the utility of these benchmarks to address broader challenges that are vital to advancing the industry.

In response to these challenges, we introduce PepPCBench, a comprehensive benchmark framework specifically designed for evaluating PFNNs on protein-peptide complex structure prediction. Using a rigorously curated and structurally diverse dataset, PepPCSet, which is temporally and structurally independent from AF3’s training data, we aim to assess the true generalizability and limitations of PFNNs in modeling peptide binding. Our evaluation includes five leading PFNNs: AF3, AFM, RoseTTAFold-All-Atom^22^ (RFAA), Chai-1^23^, and HelixFold3^24^ (HF3), across multiple dimensions including docking accuracy, confidence scoring, structural validation, and real-world applicability. By offering standardized datasets and protocols, PepPCBench serves not only as a tool for benchmarking but also as a practical guide for applying PFNNs to peptide-based drug discovery.

## 2 Results

### 2.1 Overview of PepPCBench

To evaluate the performance of AF3 in more realistic scenarios, we introduce PepPCBench, a comprehensive benchmark framework for evaluating protein-peptide complex structure prediction using PFNNs, as illustrated in **Figure 1**.

**Figure 1.**
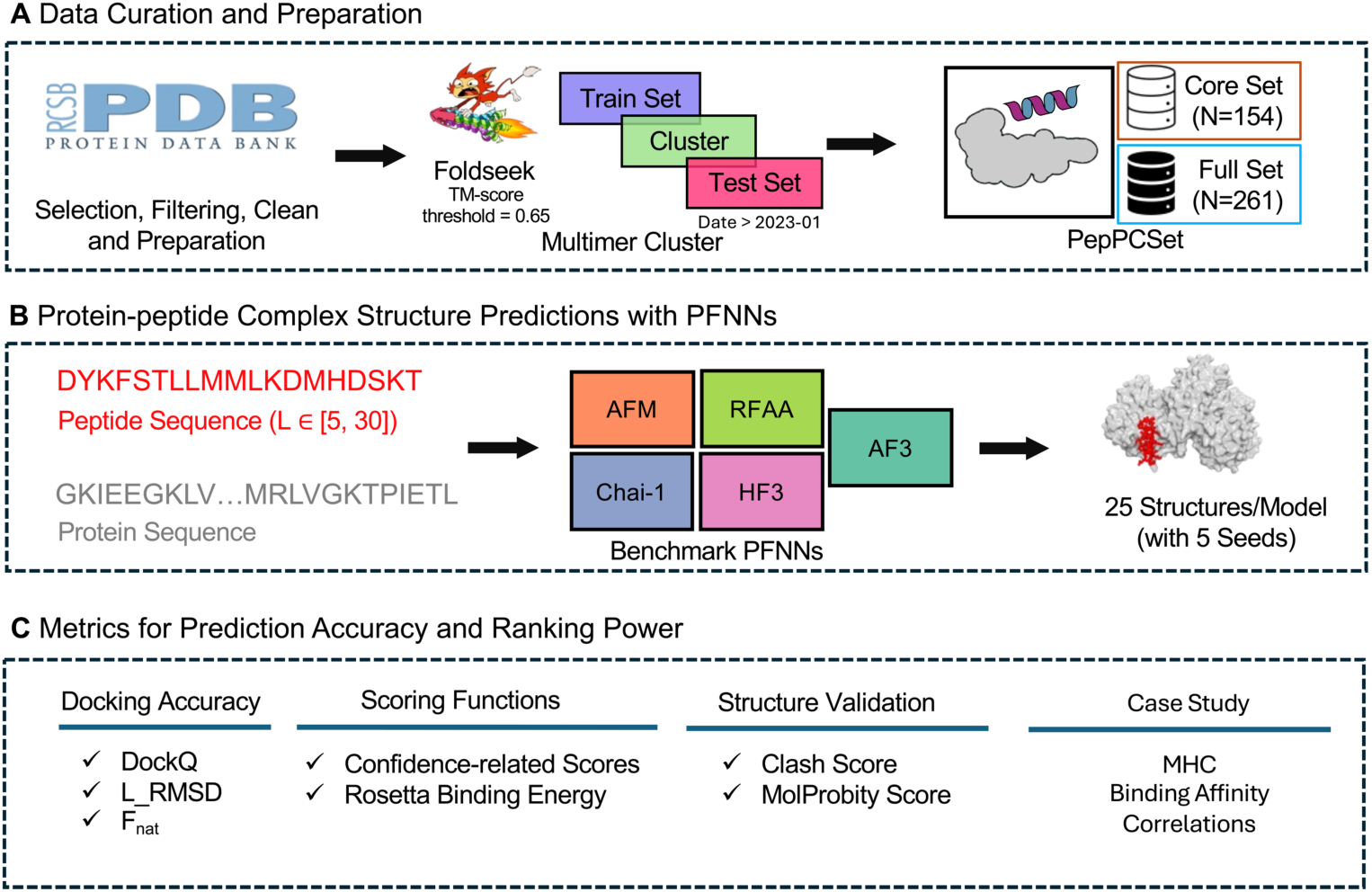
Overview of PepPCBench for Protein-peptide Complex Structure Prediction using PFNNs. (A) Data Curation and Preparation: The PepPCSet was curated from the PDB using Foldseek with a TM-score of 0.65, resulting in a “Full Set” of structures deposited from January 2023 to October 2024, and a “Core Set” by excluding similar structures from prior entries and the test set. (B) Structure Prediction: We tested PFNNs (AF3, AFM, Chai-1, HF3, RFAA) on peptides ranging from 5 to 30 residues, generating 25 models per PDB ID using five different seeds. (C) Metrics for Accuracy and Performance: Model performance was assessed using DockQ, L_RMSD, and F_nat as docking accuracy metrics, alongside model-intrinsic confidence scores and Rosetta binding energy to determine docking success rates. Structural validity was analyzed using Clash Score and MolProbity Score. We further show AF3’s docking results with experimentally validated affinity labels in a case study.

Starting with database curation, our goal was to construct a sufficiently large dataset, independent of PFNNs’ training data, to evaluate its true performance. To achieve this, we curated the PepPCSet from the PDB through a standard processing pipeline that includes selecting, filtering, and cleaning of protein-peptide complex structures. Additionally, we developed a preprocessing protocol to prepare the data for PFNN input. Next, we performed structure-based clustering using Foldseek^25^ with a TM-score^26^ threshold of 0.65. This process resulted in two datasets: the Full Set, which includes 261 complete complex structures deposited between January 2023 and October 2024, and the Core Set, which was derived by removing structures that were structurally similar to previously reported PDB entries and to other complexes within the test set, resulting in 154 non-redundant complexes. **Figure 2** illustrates the distribution of PepPCSet across three aspects: peptide length, Rosetta^27,28^ binding energy (calculated by using Rosetta Ref2015 scoring function), and the percentage of interface accessible surface area (%ASA_i) distribution. This rigorous approach ensures the reliability and diversity of the dataset for evaluation.

**Figure 2.**
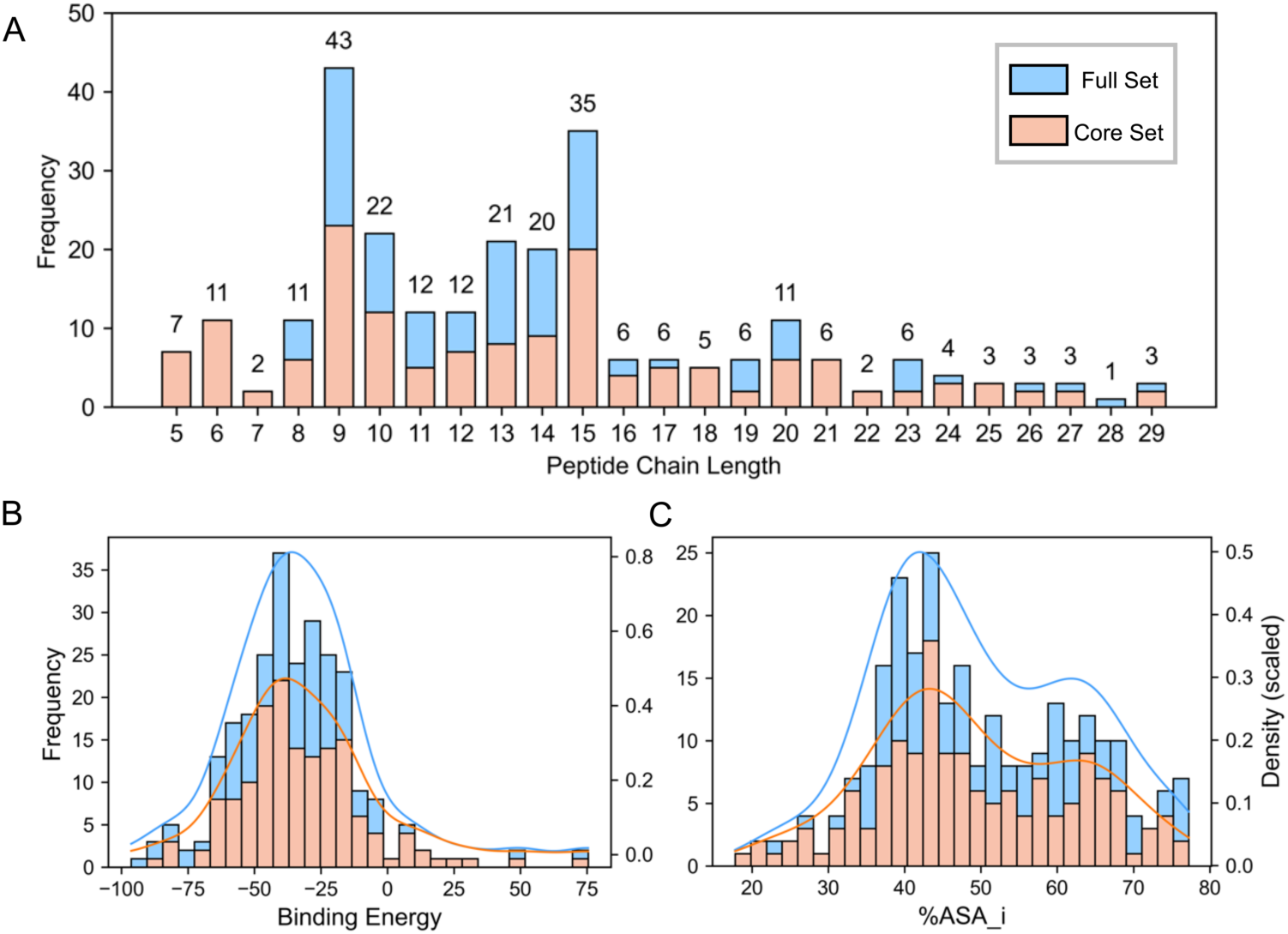
The distribution of (A) peptide chain length, (B) Rosetta binding energy and (C) %ASA_i for PepPCSet.

For protein-peptide complex structure prediction models, we evaluated five PFNNs on the PepPCSet, with peptide sequences ranging from 5 to 30 residues. For longitudinal comparison, we included AFM, and for lateral benchmarking, we assessed additional models such as RFAA, Chai-1, and HF3. To ensure robustness and reproducibility, each model generated 25 structures per complex using five distinct random seeds for further analysis.

And for metrics for prediction accuracy and ranking power: we assessed model performance across three key aspects: docking accuracy (evaluated using DockQ, L_RMSD, and F_nat), model-intrinsic confidence scores, and physics-based Rosetta binding energy scores. These metrics were used to calculate docking success rates. Additionally, we evaluated the structural validity of the predicted peptides and complexes by calculating Clash Score and MolProbity Score.^29^ Finally, we demonstrated AF3’s performance in a case study by correlating docking results with experimentally validated activity or affinity labels.

In this study, we introduced PepPCBench, a benchmark framework designed to evaluate protein-peptide complex structure prediction using PFNNs. By curating a large, independent dataset (PepPCSet), we rigorously assessed five PFNNs, including AF3, AFM, RFAA, Chai-1, and HF3, on peptides of varying lengths. We evaluated performance using docking accuracy, confidence scores, and structural validity metrics. Our results show that AF3 outperforms other PFNNs in high-quality docking predictions, with superior docking accuracy, scoring performance, and structural validity. However, our case study reveals that, despite achieving high docking precision, the application of these models for virtual screening remains a significant challenge. Through this work, we aim to drive progress in the field and help identify breakthroughs in protein-peptide complex prediction.

### 2.2 Comparison of Docking Accuracy of AF3 to Other PFNNs

In this section, we compare AF3 with other PFNNs, focusing on evaluating the accuracy of protein-peptide complex structure predictions. To effectively address the challenges in this field, we categorize the dataset into three distinct groups: (1) Similarity, which explores the impact of deep learning models on memorization issues; (2) Peptide Length, which analyzes the impact of varying peptide chain lengths on prediction accuracy; and (3) Difficulty, which is based on the difficulty levels from traditional docking methods, investigating how these levels influence the AF3 predictions. These classifications provide a more comprehensive understanding of how different factors affect the predictive capabilities of the models.

#### 2.2.1 The Effect of Structure Memorization

In the field of protein structure prediction, PFNNs are capable of inferring protein structures from amino acid sequences, identifying key features within the protein folding energy landscape. These features are defined by the diversity and frequency of various conformations. Notably, the “memorization” issue of PFNNs has become a focal point of research. Many studies related to AF have raised the question of whether AF3 truly grasps the underlying principles of protein folding energetics.^30^ This issue directly impacts our ability to accurately define the limits of its predictive capabilities, particularly when it comes to the effectiveness of predicting rare proteins. Therefore, further validation of AF3’s prediction power, as well as exploration of physics-based assisted methods, has become crucial.

Recent studies have shown that AF3’s exceptional performance is not entirely attributable to its understanding of protein energetics. Chakravarty et al.^31,32^, by testing AF3’s predictive limits on folding transition proteins, found that some of its successful predictions actually stem from “memorizing” structures in the training set, rather than relying on learned energetic principles. Similarly, Škrinjar et al.^33^, by constructing a dataset called “Runs N’ Poses,” studied the prediction of small molecule conformations. They discovered that existing co-folding methods often depend on memorized interactions from the training data, particularly manifesting as a strong reliance on ligand poses present in the training set. This benchmark dataset includes 2600 high-resolution protein-ligand systems, whose release dates exceeded the cutoff for training these methods.

In the context of protein-peptide complex structure prediction, Zhou’s benchmark^34^ fails to adequately account for the similarity between the training data and the test set, leading to an overestimation of performance. Manshour et al.^18^, recognizing the training cutoff date of AF3, applied a sequence-based CD-HIT^35^ method to further reduce redundancy, thereby providing a more objective evaluation of AF series’ performance. However, their study still faces limitations due to the relatively small size of the dataset and potential concerns about the scientific rigor of their dataset splitting approach. Consequently, there is a need for further refinement in dataset construction and testing methodologies to better assess AF3’s capabilities.

Here, we built a dataset called PepPCSet to test AF3’s ability in protein-peptide complex structure prediction. To reduce redundancy and ensure data independence, we clustered the dataset using the Foldseek algorithm with a TMScore threshold of 0.65. This allowed us to divide the Full Set dataset into two subsets: Train (N=77) and Test (N=184). The Train Set consists of portions of the Full Set that have a similarity greater than the threshold with the Train structures, while the remaining data is allocated to the Test Set. The Core Set, on the other hand, eliminates redundancy within itself, containing 154 unique structures, providing a more reliable dataset for validating AF3’s performance in predicting peptide complexes.

As shown in **Figure 3**, we compared the Full, Core, Train, and Test datasets. The results revealed that AF3 showed significant differences in DockQ scores between the Train (0.701) and Core (0.458), Test (0.465) datasets, indicating a strong “memorization” effect in AF3’s structure predictions. A similar phenomenon was observed with other PFNN methods, suggesting that the “memorization” problem is widespread and has a notable impact. Therefore, when using AF3 for structure prediction, users should carefully consider this issue. To improve prediction accuracy, we recommend performing an initial structural similarity calculation of the AF3 training set using tools like Foldseek before prediction and using this information to adjust the confidence level of the prediction results, thus mitigating the influence of the “memorization effect” on the final predictions.

**Figure 3.**
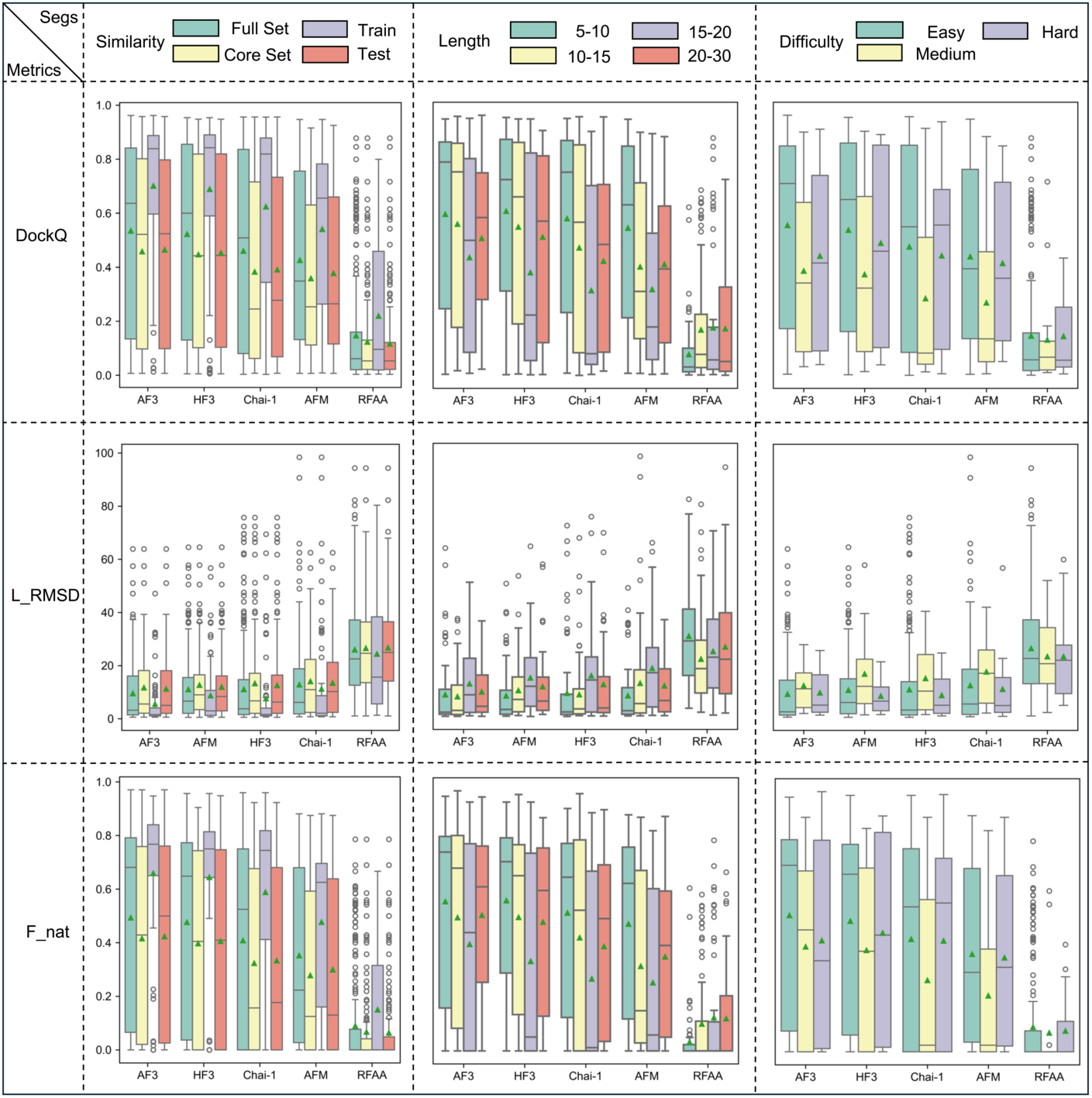
Comparison of docking accuracy between AF3 and other PFNNs. This figure presents three metrics (rows: DockQ, L_RMSD and F_nat) across three classifications (columns: length, difficulty and similarity) evaluated for five PFNNs. The data points for each metric represent the top-1 value for each PDB ID. The *x-axis* labels are ordered according to the mean value of each corresponding metric.

#### 2.2.2 The Effect of Peptide Length

Peptide chain length is a key factor influencing docking accuracy. To further explore the impact of peptide chain length on the binding prediction accuracy of protein-peptide complexes, we divided the entire dataset into four subsets based on peptide chain length, as shown in the length distribution in **Figure 2A**. Each interval contains a relatively balanced number of peptides, organized using a segment strategy. The effect of peptide chain length on docking accuracy is illustrated in **Figure 3** (Column 2). The results indicate that when the peptide chain length is less than 20, PFNNs’ DockQ and F_nat scores decrease as the peptide chain length increases, while the L_RMSD score increases (the reasons for the low accuracy of RFAA will be further discussed in the Methods section). When the peptide chain length is between 5-10, AF3 achieved the highest DockQ score of 0.597, slightly below HF3’s score of 0.601. However, in the peptide length range of 15-20, AF3 significantly outperformed HF3, with scores of 0.437 and 0.381, respectively. Therefore, AF3 performs excellently in the longer peptide chain groups and maintains a leading position in the overall average ranking.

The decrease in prediction accuracy of PFNNs with increasing peptide chain length may be related to several factors, particularly the structural flexibility of the peptide chain. Longer peptides are typically more flexible than shorter peptides and can adopt a wider range of conformations, making it difficult for the model to accurately predict their final 3D structure. Capturing all the folding details of longer peptide chains may exceed the model’s capacity. Moreover, these metrics are compared against PDB crystal structures, which represent only one of many possible conformations, thus failing to comprehensively reflect all potential states. As the peptide chain length increases, the interaction sites with the protein may become more complex and variable. Longer peptides are likely to interact with multiple locations on the protein, making it challenging for the model to capture all the key interaction sites, especially when there is insufficient experimental data for training.

This finding aligns with traditional docking models, which typically show decreased docking accuracy as peptide chain length increases.^16^ However, we observed an improvement in PFNN performance when the peptide chain length exceeded 20. A peptide length of 20 amino acids seems to be a critical threshold. We further discovered that some peptides longer than 20 amino acids may share more structural features with shorter proteins. This is because protein folding tools are primarily trained on protein sequences much longer than 20 amino acids. Therefore, as the peptide chain length increases, docking accuracy improves, indicating that longer peptides may, in some cases, form more stable structures with proteins.

#### 2.2.3 The Effect of Conformational Change

In traditional peptide docking methodologies, predicting protein-peptide binding modes that undergo large conformational changes remains a significant challenge. The root mean square deviation (RMSD) of backbone atoms is a crucial quality metric for assessing conformational adaptation. This parameter quantifies the structural deviation between a peptide’s bound conformation and its idealized extended or helical reference states. Notably, elevated RMSD values are associated with increased computational demands due to the greater complexity involved in exploring the conformational space. Building on this fundamental concept, the PepSet benchmark dataset developed by Weng et al.^16^ introduces three distinct classifications of docking difficulty (Easy, Medium, Difficult) based on quantitative analysis of RMSD_bound/helical_ and RMSD_bound/extended_ ratios.

We used this classification method to obtain 222 easy, 20 medium, and 19 difficult complexes (see Methods). The prediction accuracy results are shown in Figure 3 (Column 3). The AF3 model achieved a DockQ score of 0.555 for easy complexes, 0.388 for medium, and 0.443 for difficult complexes. This trend is similar to the impact of length on prediction accuracy. Specifically, we found that all 19 difficult complexes had lengths greater than 20, while the medium-length complexes ranged from 10 to 20. When comparing only the easy and medium complexes with lengths between 10 and 20, we observed that the medium complexes performed worse than the easy ones. These findings are consistent with our dataset, confirming that the difficulty rule holds for peptides shorter than 20.

Looking ahead, models should place greater emphasis on improving the accuracy of medium-length peptides in the 10-20 range, as they represent a “gray area” in predictions. For traditional docking software, their longer length typically leads to larger conformational changes, complicating predictions. In the case of PFNNs, the lack of sufficient training data for this class of peptides may explain the lower prediction accuracy for these medium-length peptides.

### 2.3 Comparison of the Ranking Power of Confidence-Related Scores for PFNNs

The ability to rank predicted structures accurately using model-intrinsic confidence scores is crucial for real-world scenarios. In this section, we evaluate the ranking power of these confidence scores and investigate potential improvements through the incorporation of Rosetta binding energy and enhanced sampling techniques. Additionally, recognizing that physics-based scoring methods heavily rely on structural accuracy, we perform structural relaxation using Amber (modified from ColabFold’s implementation)^21^ to explore its impact alongside confidence scores on predictive performance.

Figure 4A illustrates the relationship between DockQ scores and intrinsic confidence scores. DockQ assesses structural similarity between predictions and their true counterparts, while the confidence score quantifies the model’s certainty in its predictions. Our analysis reveals strong correlations for AF3 and AFM, indicating that predictions with higher confidence scores generally achieve superior DockQ values. Notably, HF3 encountered computational errors leading to negative confidence values; to correct this, we replaced the erroneous scores with ipTM approximations.

**Figure 4.**
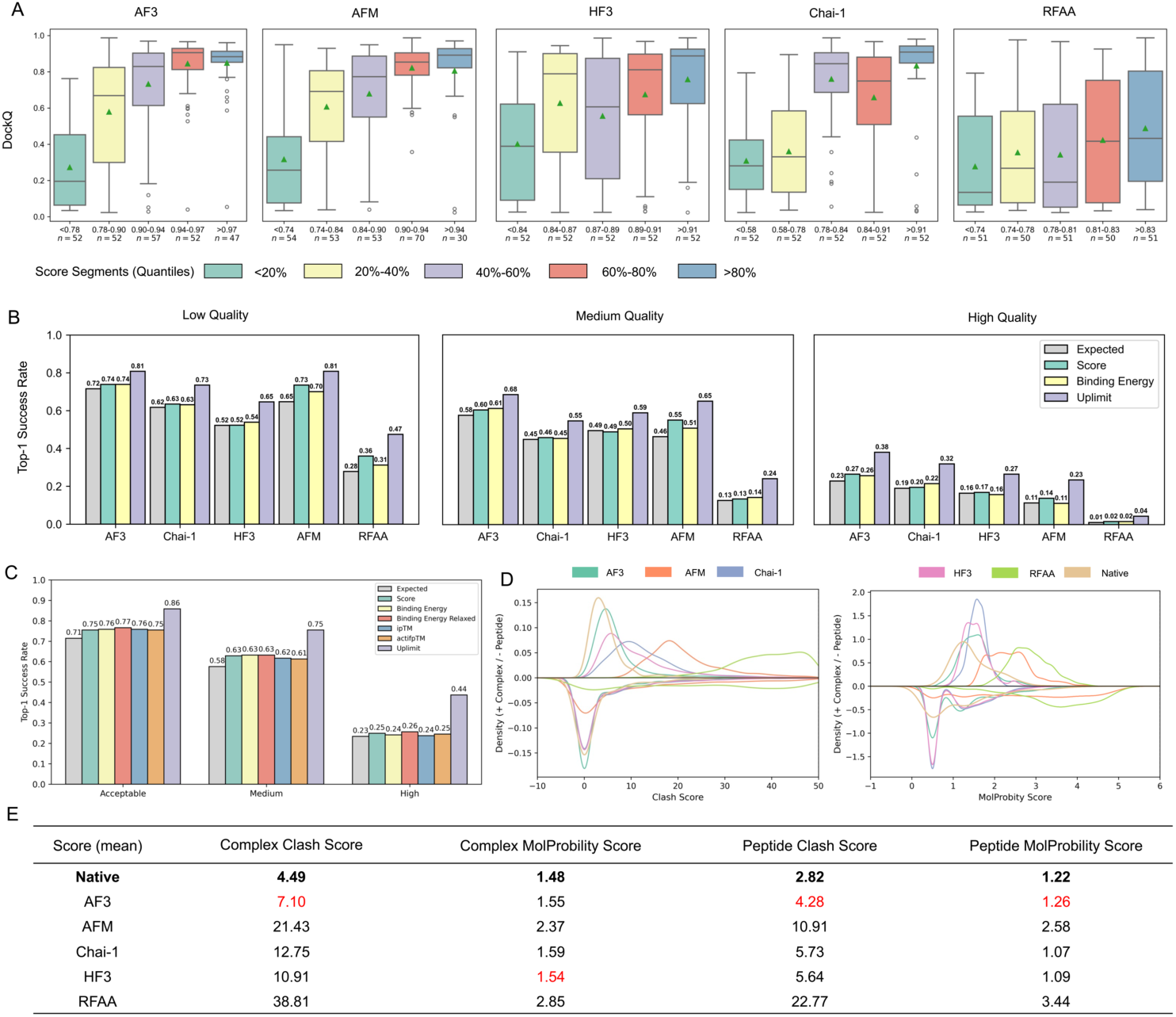
Metrics for protein-peptide complex structure prediction. (A) The relationship between DockQ and confidence-related scores of PFNNs. The scores were segmented by number quantiles, with each bin evenly dividing our test samples. (B) The top-1 success rate using confidence-related scores and Rosetta binding energy for three types (Low, Medium and High) of structure prediction quality. (C) The top-1 success rate for AF3 with enhanced sampling and relaxation for the predicted structure. (D) and (E) Distribution and statistical values for the Complex/Peptide’s Clash Score and MolProbity Score, with the native PDB score values highlighted in bold and the closest PFNN values colored in red.

To comprehensively evaluate predictive performance, we calculated the docking success rate, considering both docking quality and scoring accuracy. Docking quality was categorized into four levels—Unacceptable, Low, Medium, and High—based on F_nat and L_RMSD/i_RMSD criteria derived from CAPRI-peptide guidelines^36^ (specific thresholds provided in Methods). Figure 4B compares the performance of five PFNNs across Low, Medium, and High-quality prediction categories.

In cases categorized as Low-quality predictions, AFM’s confidence score achieved higher success rates compared to AF3. Nonetheless, AF3 exhibited greater overall robustness, reflected by its higher expected docking success rate. For Medium and High-quality predictions, AF3 consistently surpassed other models, highlighting its superior predictive capacity. Interestingly, although Chai-1 and HF3 performed worse than AFM at lower quality levels, they outperformed AFM for High-quality predictions, indicating their utility under stringent accuracy conditions.

We also investigated whether enhanced sampling could further elevate AF3’s predictive performance by increasing sampling density from 5 seeds × 5 models to 20 seeds × 10 models. Results shown in Figure 4C demonstrate that enhanced sampling minimally influenced actual predictive accuracy but notably improved the theoretical upper limit (labeled “Uplimit” in purple). Difficult cases remained challenging primarily because, despite increased sampling, we lack a more effective scoring function to distinguish correct structures in realistic, blind-prediction scenarios.

To quantify this effect, we defined two metrics: “Expected” and “Uplimit.” The “Expected” metric indicates success rates obtained by randomly selecting from predicted structures, while “Uplimit” assumes the structure classification labels were already known, representing a theoretical upper boundary. Enhanced sampling yielded a modest improvement (approximately 1–2%) in Medium-quality predictions, suggesting slight benefits for certain PDB entries; however, this gain remained limited and aligned closely with expected performance. Conversely, for High-quality predictions, enhanced sampling negatively affected performance, decreasing scoring function accuracy. This deterioration likely arose from the expanded structure pool requiring more precise scoring discrimination, thus diluting predictive accuracy to near-baseline levels.

Moreover, we examined Rosetta binding energy’s capability to discriminate correct structures relative to intrinsic confidence scores (Figure 4B). Given the dependence of physics-based scoring methods on structural validity, we relaxed AF3-generated models alongside enhanced sampling to determine if relaxation improved predictive discrimination. Relaxation outcomes mirrored non-relaxed scenarios, confirming AF3’s initial structural accuracy—a conclusion consistent with structure-validation findings in subsequent sections. Importantly, for High-quality predictions, Rosetta binding energy performed poorly without relaxation, yet outperformed intrinsic confidence scores after relaxation. Consequently, we recommend employing relaxation post-processing when using Rosetta binding energy as a scoring criterion.

This analysis evaluated the effectiveness of model-intrinsic confidence scores in ranking predicted structures and explored enhancements through improved sampling and Rosetta binding energy. We found that higher confidence scores generally correlate with better predictions, with AF3 performing particularly well in high-quality cases. Enhanced sampling had minimal impact on real-world predictions but significantly improved the theoretical success rate, highlighting the potential for further refining the scoring function to better identify native-like complex structures. Finally, Rosetta binding energy, when combined with structural relaxation, showed slight improvements over confidence scores alone.

### 2.4 The Structure Validation for Predicted Structures

MolProbity^29^ is a widely used validation tool for assessing protein structure quality by analyzing stereochemistry and atomic interactions. It evaluates amino acid residue conformations, angles, and bond lengths, identifying deviations from standard values. Manshour et al.^18^ introduced the tool for analyzing protein-peptide complex structures, providing detailed examples that highlight the differences between AF3 and AFM.

In this study, we focused on the distribution and statistical values of the structural validity for the five PFNN models, offering readers a broad understanding of their performance. As shown in Figures 4D **and 4E**, we computed the distribution and statistical values for the Clash Score and MolProbity Score of the complexes/peptides. Additionally, we present the Clash Score and MolProbity Score distributions and values for the native PDB structure for comparison. Our analysis reveals that AF3 is the closest to the native structure, as evidenced by both the distribution curves and the statistical means shown in Figure 4E. The native PDB values are highlighted in bold, and the closest PFNN values to the native ones are colored in red.

### 2.5 Case Study for AF3 Predicting Protein-peptide Complex Under Real-word Scenarios

Peptide-binding proteins play essential roles in a wide range of biological processes, and accurately predicting their binding specificity remains a long-standing challenge. Recent work by Motmaen et al.^37^ demonstrated that integrating a classification head into the AF network and jointly fine-tuning for both structure prediction and binding classification significantly improves generalization. Their model achieved competitive performance with the sequence-based NetMHCpan^38^ on MHC-I/II systems and further demonstrated strong generalization in peptide recognition by SH3 and PDZ domains, outperforming purely sequence-based approaches—especially in data-scarce scenarios.

Building on this motivation, we conducted a real-world case study to evaluate the practical utility of AF3 in virtual screening for high-affinity peptide binders in an MHC-II system. Specifically, we focused on the HLA-A*02:01 allele and curated a dataset consisting of 9-mer peptides with experimentally measured binding affinities (K_d_). To ensure a well-balanced dataset across different affinity ranges, we stratified the data into seven bins based on the logarithmic scale of the K_d_ values. From each bin, 20 peptides were randomly sampled, resulting in a representative and diverse dataset for downstream evaluation, comprising a total of 140 peptides across 7 bins.

For each peptide-MHC pair, we used various scoring functions to rank the top-1 predicted structure from AF3, which served as the proxy label for downstream analysis. As shown in Figure 5A, we first computed the Pearson correlation between scoring function outputs and the raw K_d_ values. We then repeated this analysis using log-transformed K_d_ values (Figure 5B), and observed consistently higher correlation, likely due to the non-linear nature of binding energy scales and the presence of negative values in some scoring functions.

**Figure 5.**
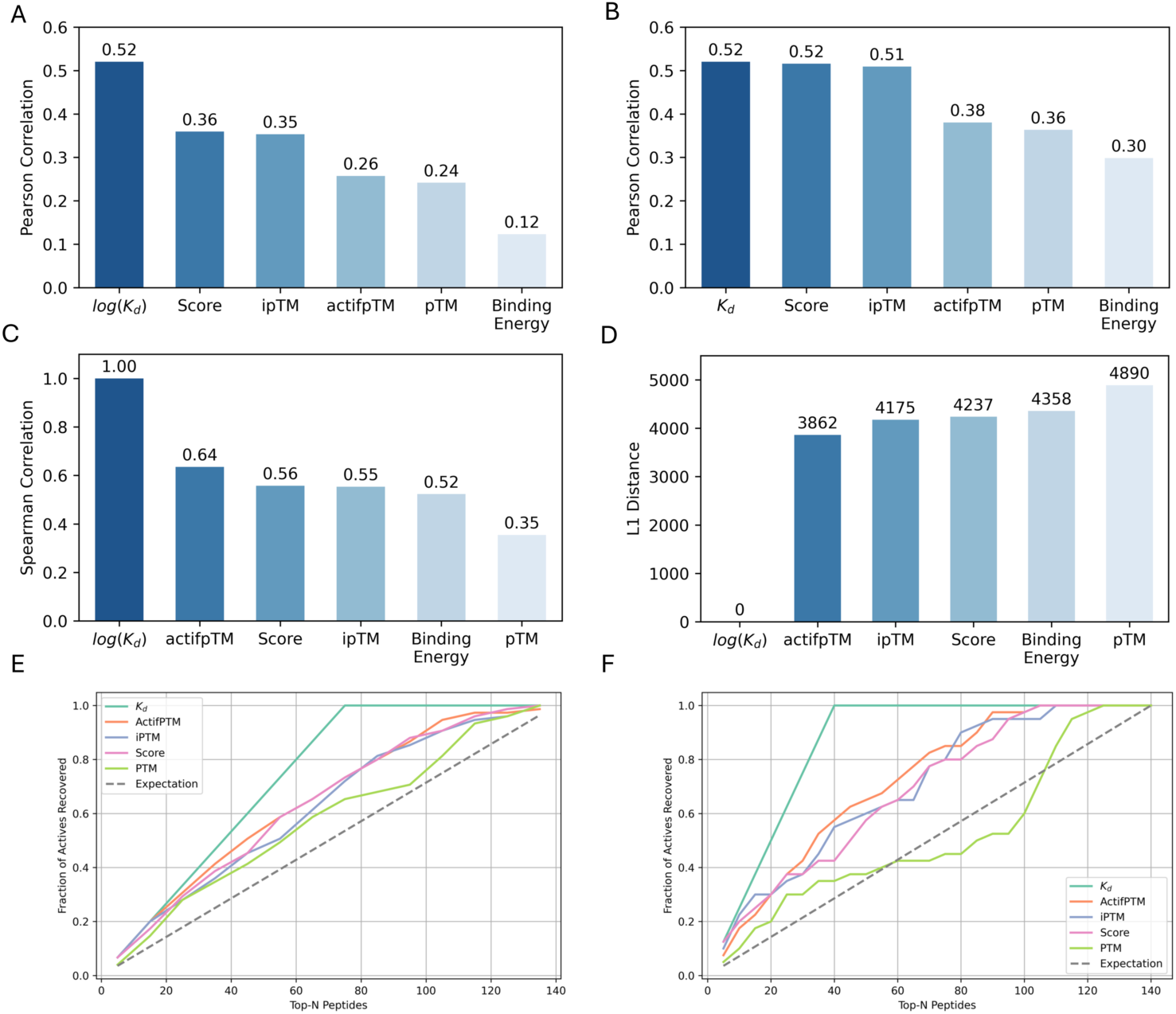
Case study of AF3 in Screening High-affinity Peptide Binders for the MHC-II System. (A) Pearson correlation of scoring function values against (A) K_d_ and (B) log(K_d_). (C) Spearman correlation of scoring function values against K_d_. (D) The L1 distance between the peptide rankings based on scoring function values and the true rankings. The active recovery rate for metrics to identify (E) active/non-active peptides with a K_d_ threshold of 500 nM, and (F) to select high-affinity peptide binders. “Score” refers to the confidence-related ranking scores output by PFNNs.

To further account for potential non-linear relationships, we calculated Spearman correlation coefficients between scores and K_d_ values (Figure 5C), which confirmed consistent trends. Beyond correlation, we evaluated how well each scoring function ranked candidate peptides by computing the L1 distance between the predicted and true rankings (Figure 5D). Among all metrics, actifpTM showed the best agreement with experimental rankings, highlighting its robustness in capturing binding-relevant structural features.

To assess the functional relevance of these scores in a screening context, we introduced the concept of active peptide recovery rate. This metric reflects the proportion of known binders recovered above a given affinity threshold. As shown in Figure 5E and **5F**, we evaluated recovery performance under two thresholds: 500 nM (active vs. non-active classification) and 10 nM (high-affinity binder selection). Interestingly, all scoring functions performed similarly in distinguishing active from inactive peptides (Figure 5E), suggesting that the baseline discrimination capability is largely determined by the quality of AF3-predicted structures. However, when the goal was to identify high-affinity binders, actifpTM exhibited slightly better performance (Figure 5F), consistent with its superior ranking behavior observed in Figure 5C and **5D**.

## 3 Discussion

In this study, we introduced PepPCBench, a comprehensive benchmark specifically designed to evaluate the performance of PFNNs, with a particular focus on AF3, in predicting protein-peptide complex structures. By leveraging a rigorously curated dataset (PepPCSet), we aimed to provide a standardized and robust platform to assess both the capabilities and limitations of current PFNNs in handling the structural complexity and flexibility inherent to protein-peptide interactions.

Our results demonstrate that AF3 outperforms its peers (AFM, RFAA, Chai-1, and HF3) in terms of docking accuracy and structural validity. It also shows superior performance in model ranking and confidence assessment, particularly when using refined confidence metrics such as actifpTM, which better account for flexible and disordered regions.

However, despite its overall superiority, AF3’s performance reveals several caveats. Most notably, our analysis confirmed that AF3 suffers from structural memorization, with inflated performance on peptide complexes similar to those seen during training. The significant drop in DockQ scores when comparing the Train Set to the Test and Core Sets underscores the importance of evaluating models on structurally novel complexes. Furthermore, peptides with intermediate lengths (10–20 residues) and moderate conformational shifts remain especially challenging, pointing to a blind spot in both AF3’s training and current data coverage. While enhanced sampling improved the theoretical success rate, the limited effectiveness of current scoring functions hindered meaningful gains in actual predictive performance.

Beyond docking metrics, we validated structure quality using MolProbity and Clash Score, finding that AF3-generated models are not only geometrically sound but also closely approximate native PDB structures. This structural fidelity lays a strong foundation for downstream functional predictions, such as virtual screening.

Indeed, our case study on MHC-II systems demonstrated AF3’s promising application potential in real-world settings. Using experimentally measured binding affinities, we observed that AF3’s structural predictions—when combined with robust scoring functions like actifpTM and Rosetta binding energy—can reliably rank peptide binders and recover high-affinity ligands. However, ranking performance varied depending on the scoring function used, reinforcing the importance of combining structural prediction with reliable post hoc evaluation.

PepPCBench addresses a critical need in the field by offering a rigorous, diverse, and reproducible benchmark for evaluating PFNN-based protein–peptide complex prediction. Through our assessment of five state-of-the-art models, we uncovered key strengths and limitations—particularly those of AF3—highlighting its promise for structure-based peptide design, while also revealing areas for improvement. Our findings emphasize the importance of methodological transparency, dataset independence, and the integration of complementary validation strategies in future model development.

While PepPCBench establishes a strong foundation, it also points to clear avenues for enhancement—such as incorporating more diverse protein families, post-translational modifications, and cyclic peptides to improve benchmark generalizability. We believe PepPCBench will be a valuable resource for both method developers and experimental biologists, accelerating the advancement of next-generation peptide therapeutics through more reliable structure prediction.

## 4 Methods

### 4.1 Data Curation and Preparation

In this study, we collected protein-peptide complex structures from the RCSB PDB database to evaluate model performance. The dataset included structures released between January 1, 2023, and October 10, 2024. To ensure a high-quality dataset and precise evaluation, we applied a rigorous selection and standardization process as follows:

Initial Selection Criteria:

1. Entries were restricted to those labeled as “prot” to exclude structures containing nucleic acids.
2. Each structure included at least two chains, with one being a peptide chain containing more than 5 and up to 30 residues.
3. Only structures determined by NMR, X-ray diffraction, or electron microscopy with a resolution of ≤ 3.0 Å were considered.

Standardization Process:

1. Peptides had to form a continuous chain through the backbone atoms (N, CA, C, O).
2. Structures with missing backbone atoms (N, CA, C, O) in any protein residue were removed.
3. Proteins were required to contain at least 50 residues or be at least three times the size of the peptide chain to ensure a sufficient binding interface.

Content-Based Filtering:

Structures were further filtered by examining the “Structure Title” for keywords such as “complex with” or “bound to” to confirm the presence of a protein-peptide interaction.

This multi-step process ensured the construction of a high-quality dataset suitable for evaluating model performance under stringent conditions.

#### 4.1.1 Data Preparation and Structure Clustering

The sequences for the receptor and peptide to be modeled were obtained from the SEQRES lines in the PDB files, which reflect the expressed constructs rather than the resolved structures. Unknown residues were removed, and non-canonical amino acids (NCAA) were converted to canonical ones based on AF3’s CCD dictionary, ensuring smooth operation of the model. Missing heavy atoms were added using pdbfixer, incorrect chain labels were corrected with pdbtools, and chain splitting was performed using PyMOL. These processed files are available on our GitHub.

The dataset was clustered based on structural similarity using the Foldseek tool, with the easy-multimercluster module designed for multimer-level clustering. We applied this tool to cluster our self-built PepPCSet. First, we queried all protein-peptide complex structures with the context constraint “peptide” prior to the data cutoff of 2021-09-30 and collected the corresponding mmCIF files for the next clustering step. After mixing the collected training data and PepPCSet, we clustered the dataset and selected the highest-resolution structure from each cluster as the representative entry for the final dataset.

To reduce redundancy and ensure data independence, we applied the Foldseek algorithm with a TMScore threshold of 0.65. This allowed us to divide the Full Set dataset into two subsets: Train (N=77) and Test (N=184). The Train Set consists of structures from the Full Set that have similarity greater than the threshold with the Train structures, while the remaining data was assigned to the Test Set. The Core Set eliminates redundancy, containing 154 unique structures, and provides a more reliable dataset for validating AF3’s performance in predicting peptide complexes.

### 4.2 Protein Folding Neural Networks

With the rapid expansion of PFNNs, each method brings its own unique strengths, as summarized in Table S1. In this study, we compared AF3 against four different PFNNs to evaluate their relative performance.

#### 4.2.1 AF3

Modeling was conducted using the publicly accessible AF3 repository (https://github.com/google-deepmind/alphafold3). We utilized the local standalone version (Docker container) for batch processing the prediction and analysis pipeline. No additional refinement was performed on the models. The MSA configuration adhered to the original default settings, and we diligently sourced MSA and template information for each input peptide, ensuring all MSA and template data were cut off before September 30, 2021, to prevent training data leakage. A single A800 GPU (80G) was used for the inference step, and Python (version 3.11) scripts were employed for process management and result analysis.

#### 4.2.2 AFM

AFM was introduced in 2022, and since then, several tools have been developed to simplify its use. One standout tool is ColabFold^21^ (CF), which enhances the accessibility and speed of AF and AFM through Google Colab. By streamlining the workflow, CF has significantly accelerated protein complex predictions, making protein structure modeling faster and more user-friendly. This advancement has greatly contributed to progress in structural biology, democratizing access to protein structure predictions and facilitating research in the field.

To create a local version, we utilized a tool called localcolabfold for fast MSA/template searching and inference. This tool enables efficient processing on local machines, streamlining the workflow for protein structure prediction while maintaining the speed and convenience of the ColabFold interface.

#### 4.2.3 RFAA

RoseTTAFold All-Atom (RFAA) is an advanced neural network designed for generalized biomolecular modeling and design. It extends the capabilities of previous models like AlphaFold2 and RoseTTAFold by incorporating the ability to model full biological assemblies. This includes proteins, nucleic acids, small molecules, metals, and covalent modifications based on the sequences of polymers and the atomic bonded geometry of the small molecules and covalent modifications.

In this study, we utilized the GitHub version of the tool and encountered scalability issues with peptide lengths shorter than 10 amino acids, as the program does not effectively handle these shorter peptides. Additionally, the 3D atomic positions generated by RFAA exhibited numerous conflicts and clashes, both within peptide chains and between peptide and protein chains (also failed on protein-protein interactions). These two issues significantly impacted the performance of RFAA in our analysis, leading to suboptimal results.

#### 4.2.4 Chai-1

Chai-1 is a multi-modal foundation model designed for molecular structure prediction, demonstrating state-of-the-art performance across a variety of tasks crucial for drug discovery. It’s distinguished by its ability to incorporate experimental restraints, such as data derived from wet-lab experiments, which significantly enhances its prediction capabilities. Chai-1 can operate effectively both with and without multiple sequence alignments (MSAs), maintaining high performance in single-sequence mode.

In this paper we use the local version provided at https://github.com/chaidiscovery/chai-lab.git.

#### 4.2.5 HelixFold3

HelixFold3^24^ is a biomolecular structure prediction model developed by Baidu’s PaddleHelix team, released on January 15, 2025. It aims to replicate the capabilities of DeepMind’s AlphaFold3, focusing on predicting the 3D structures of proteins, RNA, DNA, small molecules, and their complexes.

### 4.3 MSA

Multiple Sequence Alignment (MSA) is crucial in protein structure prediction, helping to infer 3D structures and functions by identifying conserved regions linked to structural stability and biological activity. MSA also reveals co-evolutionary signals, which are valuable for predicting contact maps and improving structural models, with techniques like Mutual Information (MI) and Direct Coupling Analysis (DCA) enhancing accuracy. In homology modeling, MSA aligns target proteins with templates, aiding precise predictions. While deep learning models like AF and RoseTTAFold rely on MSA to boost performance, challenges such as low sequence similarity and limited homologous sequences can affect alignment quality. In such cases, single-sequence methods like ESMFold may be alternatives. Despite AlphaFold’s reduced MSA dependence, high-quality MSA remains key to improving structural prediction.

The protein-peptide complex structure prediction was performed using AF3^14^. For the genetic searching step, we followed the official AF3 guidelines to ensure the optimal multiple sequence alignment (MSA) generation process. Detailed MSA parameters, including database and search configurations, are available at https://github.com/google-deepmind/alphafold3 (accessed on 2024/12/27). Here, we briefly describe the databases used for protein chain searches. Five databases were utilized: UniRef90^39^, UniProt^40^, Reduced BFD^41^, MGnify^42^, and Uniclust30^43^ + BFD^44^. The searches were performed using the Jackhmmer^45^ and HHBlits^46^ tools. During the model inference step, we fixed the random seed to ensure reproducibility of the predicted complex structures. Five models were generated in total, and the highest-ranked model was selected for downstream analysis based on AF3’s well-calibrated confidence measures, which align closely with prediction accuracy.

The detailed MSA configuration can be found in SI.

### 4.4 Evaluation metrics

#### 4.4.1 DockQ with F_nat, L_RMSD and i_RMSD

DockQ^47^ is a continuous quality metric for protein-protein docking, combining fraction of native contacts (F_nat), ligand root-mean-square-deviation (L_RMSD), and interface RMSD (i_RMSD) into a single score within [0, 1] to evaluate docking model quality. These three metrics, standardized by CAPRI^48^, assess different aspects of model accuracy and should be considered together. While CAPRI classifies models into four categories, DockQ provides a higher-resolution, quantitative measure, allowing for more precise evaluations using Z-scores or top-ranked model sums. This method has been invaluable in the CASP community and is essential for developing scoring functions in protein docking. When applied to CAPRI models, DockQ closely replicates the original classification with 94% PPV and 90% recall, eliminating the need for ad-hoc thresholds.

In this study, we employ DockQ to quantitatively evaluate the quality of protein-peptide docking results. Although DockQ was originally developed for protein-protein multimers, its application to peptide complexes requires careful consideration. Marcu et al.^36^ demonstrated that relaxing the backbone (and ligand) RMSD threshold, while imposing a restriction on side-chain RMSD, improves the selection of high-accuracy models. Fortunately, DockQ v2 has enhanced its capabilities, offering automatic quality assessment for protein multimers, nucleic acids, and small molecules, and now includes functionality for protein-peptide complexes.^49^ By simply adding the “--capri_peptide” flag, DockQ can assess protein-peptide docking models.

For automated alignment of chains, we implemented a fuzzy matching algorithm for maximum chain correlation, followed by manual correction of any erroneous chain mappings. To further streamline the pipeline, we initially rename the original peptide chain in the complex (both nature and model structures) as chain A, and group the remaining chains together as chain B.

RMSD is the primary metric used to evaluate the accuracy of model docking, encompassing several subcategories. While DockQ offers i_RMSD and L_RMSD, we have also implemented backbone and all-atom RMSDs for more detailed analysis of docking accuracy. RMSD values for the individual peptide and receptor structures were calculated using Python scripts, utilizing a data schema adapted from AF3.

#### 4.5.4 The Docking Quality Classification and Success Rate

In this study, we use the CAPRI success criteria^36,50,51^ for protein-peptide complex structure predictions:

Measures:

1. Interface residues: < 8.0 Å, between any two CB atoms (CA for Gly) across interface.
2. Native contacts: < 4.0 Å, between any two atoms across the interface (residue-based)

Classification:

1. High quality: F_nat [0.8, 1.0] and (L_RMSD < 1.0 or I_RMSD < 0.5).
2. Medium quality: F_nat [0.5, 0.8] and (L_RMSD < 2.0 or I_RMSD < 1.0) or F_nat [0.8, 1.0] and (L_RMSD > 1.0 and I_RMSD > 0.5)
3. Low quality: F_nat [0.2, 0.5] and (L_RMSD < 4.0 or I_RMSD < 2.0) or F_nat [0.5, 1.0] and (L_RMSD > 2.0 and I_RMSD > 1.0)
4. Unacceptable: The rest

We also introduced a loosen success criterion for evaluating protein-peptide complex structure predictions, adapted from PepSet^16^. The criteria are summarized as follows and the results shown in SI:

1. Low-quality (Near-native prediction): F_nat > 0.2 and L_RMSD < 7 Å.
2. Medium-quality: F_nat > 0.5 and L_RMSD < 5 Å.
3. High-quality: F_nat > 0.7 and L_RMSD < 3 Å.

We use ranking scores to select the top-1 predicted structure and determine whether it meets the quality thresholds. If it does, we label the corresponding PDB ID as a success. The success rate is then calculated as the ratio of successful PDB IDs to the total number of entries in a given dataset.

Here, we also explain how the Expected success rate is calculated. This rate represents the proportion of successful predictions across all structures, simulating the probability that the model can correctly identify structures meeting the quality threshold under random guessing conditions. This metric helps us understand the lower bound for different PFNNs and the scoring functions used.

#### 4.5.3 %ASA_i

For each protein-peptide complex, the solvent-accessible surface areas (SASA) of the protein (S1), peptide (S2), and the entire complex (S3) were calculated using freesasa^52^ (version 2.1.2). The interface area between the protein and peptide (S4) was then determined using the formula: S4 = (S1 + S2 − S3)/2, and the percentage of interface accessible surface area (%ASA_i) was defined as S4/S2 × 100%.

#### 4.5.2 Clash Score and MolProbity Score

For structure validation, we use MolProbity^29^ Score and Clash Score to complement the DockQ metrics, addressing the limitations of the latter. MolProbity is a robust structure-validation tool that assesses model quality both globally and locally, applicable to proteins and nucleic acids. It identifies and corrects structural errors through optimized hydrogen placement, all-atom contact analysis, and updated geometric criteria. With its automatic correction capabilities and intuitive graphical outputs, MolProbity significantly improves the accuracy and reliability of structural refinement. In this work, we use its Docker container version for the MolProbity web tool (which calculates both scores) to streamline batch processing, with further details provided in the *Data and Code Availability* section.

#### 4.5.3 Difficulty

Follow the work of pepATTRACT^53^ and PepSet^16^, three idealized conformations for each peptide were generated using the Python library PeptideBuilder^54^. The backbone dihedral angles and conformation types were set as follows.

1. Helical conformation: φ = −57°, ψ = −47°.
2. Extended conformation: φ = −140°, ψ = 130°.
3. Polyproline II conformation: φ = −78°, ψ = 149°.

The difficulty levels were assigned according to the following criteria, leading to the categorization of complexes in the dataset into 222 easy, 20 medium, and 19 difficult cases:

1. Easy: RMSD_bound/helical_ or RMSD_bound/extended_ ≤ 4 Å.
2. Medium: 4 Å < (RMSD_bound/extended_ and RMSD_bound/helical_) ≤8 Å.
3. Difficult: (RMSD_bound/helical_ > 8 Å and RMSD_bound/extended_ > 4 Å) or (RMSD_bound/helical_ > 4 Å and RMSD_bound/extended_ > 8 Å).

### 4.6 Scoring Functions

#### 4.6.1 Confidence-related Scores

In protein structure prediction, confidence scores are essential for assessing the reliability of predicted structures. Two common confidence scores used in models like AlphaFold3 (AF3) are pTM (predicted Template Modeling score) and ipTM (predicted Interactions with Protein Molecules). These scores are derived from the Template Modeling (TM) score, which measures the overall accuracy of the structure and is less sensitive to localized inaccuracies.

1. pTM evaluates how accurately the overall structure of the complex is predicted by comparing it to a hypothetical true structure. A pTM score above 0.5 suggests that the predicted fold is similar to the true structure, while a score below 0.5 indicates a likely incorrect prediction. However, the pTM score can be influenced by the accuracy of larger subunits, potentially overshadowing errors in smaller subunits.
2. ipTM focuses on the predicted relative positions of the subunits in a protein-protein complex. Scores above 0.8 indicate high confidence, while scores below 0.6 suggest a likely failure. Scores between 0.6 and 0.8 are considered ambiguous. In large-scale screenings of protein-protein interactions, lower ipTM thresholds (e.g., 0.3) are sometimes used for initial screening, with further analysis for higher-scoring pairs.

While both scores are important, ipTM may be more useful because the accuracy of the subunit positions directly impacts the overall structure prediction. Therefore, confidence in multimers should be based on a combination of metrics, including pTM, ipTM, pLDDT, and PAE.

To improve AlphaFold’s performance for protein complexes, particularly for flexible or disordered regions, the actifpTM metric was introduced.^55^ This modification addresses flaws in the original ipTM score, which averages confidence across all interchain residue pairs, including flexible or poorly resolved regions, resulting in misleading confidence scores. actifpTM isolates the interface residues, focusing only on high-confidence regions and excluding low-confidence flexible areas. This ensures the metric more accurately reflects the quality of the interface, especially for peptides or disordered regions.

Additionally, the ranking score in AF3 is defined as a weighted combination of confidence metrics: confidence score = 0.8 × ipTM + 0.2 × pTM. In addition to this score, structural quality factors such as disorder regions and atomic clashes in the predicted structures are also taken into account to better assess and rank the reliability of the predictions.

#### 4.6.2 Rosetta Binding Energy

Rosetta binding energy is a key metric computed using the Rosetta molecular modeling software. It is used to evaluate the binding strength and stability between two molecules—such as protein-ligand, protein-peptide, or protein-protein complexes. This energy metric is widely applied in docking predictions, mutation screening, and protein engineering to assess binding affinity. In this study, we compute the Rosetta Binding Energy using the PyRosetta interface, enabling flexible and scriptable energy evaluation within a Python environment.

## Data and code availability

The data, code, and other resources necessary to reproduce the results, along with documentation and tutorials, are available on GitHub at https://github.com/zhaisilong/PepPCBench.

## CRediT authorship contribution statement

Silong Zhai and Huifeng Zhao: Writing – review & editing, Writing – original draft, Methodology, Investigation, Conceptualization. Jike Wang, Shaolong Lin and Tiantao Liu: Writing – original draft. Dejun Jiang, Huanxiang Liu and Yu Kang: Writing – review & editing. Xiaojun Yao: Writing – review & editing, Supervision, Project administration, Funding acquisition. Tingjun Hou: Writing – review & editing, Supervision, Resources, Project administration, Conceptualization.

## Acknowledgements

This work was financially supported by National Key R&D Program of China (2024YFA1307501), Natural Science Foundation of Zhejiang Province of China (LD22H300001), Science and Technology Development Fund, Macau, SAR (No. 0030/2024/RIA1) and Macao Polytechnic University (No. RP/FCA-15/2023). The manuscript was approved by Macao Polytechnic University with the submission code s/c fca.54cd.d123.b.

